# A multi-tissue transcriptome strategy for genomic resource development in *Pinus edulis*

**DOI:** 10.64898/2026.10.05.756810

**Authors:** Diana S. Macias, Ethan F. Gyllenhaal, Joseph A. Kleinkopf, William T. Pockman, Hannah E. Marx

## Abstract

**Premise:** Conifers comprise one-sixth of global forests and landscape genomic studies are urgently needed to understand their biology, yet relatively few conifer species have reference genomes. Targeted exon-capture is a practical alternative to whole-genome sequencing, but selecting loci without a reference genome remains a barrier. We developed a multi-tissue reference transcriptome for piñon pine (*Pinus edulis*) to ask how multi-tissue sampling improves probe design, and what is lost under a conventional single-tissue approach.

**Methods:** We assembled a reference transcriptome from four diploid tissues (needle, phloem, root, vegetative bud) and one haploid tissue (megagametophyte), using the latter as a pseudo-guide. We designed a target-capture probe-set by prioritizing highly expressed *P. edulis* transcripts and transcripts matching a conifer exon database.

**Results:** We recovered 76% of conserved plant genes with few duplicated transcripts. Filtering yielded 15,664 transcripts totaling 31.4 Mb, ∼0.1% of the genome. Tissue-specific expression was substantial: 27.4% of transcripts occurred exclusively in non-needle tissues. Functional annotation confirmed complementary contributions, including seed-desiccation and germination genes from megagametophytes, and defense- and stress-associated genes from roots and phloem.

**Discussion:** We built a transcriptome and efficient probe set using a roadmap suitable for other non-model plants with large genomes; keys to success were prioritizing complementary tissues and using expression-based filtering.

## 1. INTRODUCTION

Understanding the genomic basis of trait variation, and how genotype and phenotype vary across a species’ range is essential to predict responses to climate change and assess the adaptive potential of populations (Flanagan et al., 2018). This is particularly important for widespread, ecologically important tree species like piñon pine (*Pinus edulis* Engelm.), which is undergoing climate-change driven mass mortality, range shifts, and altered recruitment dynamics (Shriver et al., 2022). Pre-genomic studies of piñon pine suggest that this variation has a genetic basis. For example, individual variation in resistance to herbivory and environmental stress had a genetic basis among piñon pine populations in Arizona (Mopper et al., 1991). Similarly, a single gene variant was found to be strongly associated with performance under variable soil moisture (Mitton and Duran, 2004). If rapid adaptation in natural populations is predominately fueled by standing genetic variation (Ahrens et al., 2019), then trees and populations with key genetic variants that favor survival during drought will play a critical role. Similarly, neighboring populations of *Pinus* taxa may play a pivotal role through introgression of adaptive alleles and their associated traits (Marques et al., 2019; Buck et al., 2023). Realizing this potential for management and climate-change mitigation (Aitken and Whitlock, 2013) requires genomic resources that piñon pine has historically lacked.

Until recently, genomic studies have been limited by the size and complexity of tree genomes, especially those of conifers, whose genomes span 8-34 Gigabase pairs (Gb) and make assembly and bioinformatics challenging (De La Torre et al., 2014; Isabel et al., 2020). Despite these challenges, advancements in forest tree genomics have improved our understanding of the genetic basis of complex traits, signatures of adaptation to climate, and estimating adaptive potential (Neale and Kremer, 2011; Plomion et al., 2016; Feng and Du, 2022). One approach to characterize the distribution and evolution of adaptive diversity is to undertake reduced representation approaches that capture a consistent subset of genomic regions. There are two general approaches to genome partitioning. The first approach is to use partitioning methods that search for anonymous markers (i.e., RADSeq) which can identify and score thousands of genetic markers randomly distributed across the target genome using restriction enzymes (Davey and Blaxter, 2010). However, this process does not specifically target functional regions, and is difficult to replicate across studies due to variation in size-selection protocols (Parchman et al., 2018). The second method, exon capture, achieves genome partitioning through selective enrichment of preselected regions of the genome (Jones and Good, 2016). This approach can target conserved regions across taxa (e.g., UCEs, Faircloth et al., 2012) or regions of known function within a given taxon (e.g., exon capture, Hodges et al., 2007). The benefit of target capture is that it provides potentially deeper coverage and more accurate single nucleotide polymorphisms (SNP) calling at these focal regions, which in turn makes large sequencing projects of non-model organisms economically feasible (Gnirke et al., 2009; Hodges et al., 2009). The challenge is that designing targeted capture probes for preselected genomic regions of interest requires prior sequence knowledge, a difficult task for taxa that lack high-quality genomic resources (Jones and Good, 2016).

Transcriptomics is a method that quantifies and sequences all expressed RNA transcripts, providing a catalog of expressed genes suitable for the design of exon capture probes *de novo* to profile genomic regions of interest (Canales et al., 2014; Li et al., 2015; López de Heredia and Vázquez-Poletti, 2016; Syring et al., 2016; Baker et al., 2018) and a versatile tool for functional genomics (Ward et al., 2012). Although transcriptome sequences contain less information than whole genome sequences, they are enriched for functional information. By sequencing whole RNA from multiple developmental stages, tissue types, or treatments, one can build a draft transcriptome that catalogs the functional genes that are expressed on a regular basis across an organism’s lifetime. These different sample types can provide distinct sets of information that reveal variation in gene regulation, transcription, alternative splicing, and tissue specific function. Expression studies in *Pinus* have separately profiled embryogenesis (Aquea and Arce-Johnson, 2008), root development (Brinker et al., 2004), xylem and secondary cell wall formation (Egertsdotter et al., 2004; Bomal et al., 2008; Paiva et al., 2008; Yang et al., 2017), response to pathogenic fungi (Myburg et al., 2006; Adomas et al., 2007; Heller et al., 2008), and cold acclimation (Heath et al., 2002; Joosen et al., 2006)—underscoring how much of the expressed gene space is tissue- and stage-dependent. *De novo* conifer transcriptomes not only benefits from incorporating the diversity across developmental stages and tissues (López de Heredia and Vázquez-Poletti, 2016; Cervantes et al., 2021), but it can also leverage reads from haploid megagametophyte tissue as a guide for assembly to substantially increase transcriptome completeness and allow the detection of paralogs (Ojeda et al., 2019).

Here, we developed two paired genomic resources for piñon pine: a multi-tissue *de novo* transcriptome and an associated set of exon capture probes. Our study began in the absence of a *Pinus edulis* reference genome, where the most closely related genomic resource was a highly fragmented draft genome (4.2 million scaffolds) of sugar pine *(Pinus lambertiana* Dougl.) (Stevens et al., 2016), a species that diverged from pinon pine ∼60 million years ago (Jin et al., 2021). Additionally, we aimed to capture a diverse set of genes and expression profiles by sequencing transcriptomes from different tissues and developmental stages, including the bioinformatically helpful haploid megagametophyte stage (Cervantes et al., 2021). The specific motivation for our exon-capture probe design was to conduct a landscape genomic study that would pinpoint geographic regions and allelic variation that may provide resilience against climate change. We sought to build a more comprehensive transcriptome reference, and in turn, a target capture probe set more likely to include genes underlying climate related functions such as water retention, resistance to fungal pathogens, and production of insect-resistant secondary compounds. In doing so, we asked a question central to designing capture probes without a reference genome: how much does multi-tissue sampling contribute to a probe set, and what would a conventional single-tissue approach miss? By quantifying the targets lost under needle-only sampling, we evaluate tissue breadth as an explicit design variable in transcriptome-derived target capture.

## 2. METHODS

### 2.1 Sampling and sequencing

#### 2.1.1 Study species

Piñon pine (*Pinus edulis* Engelm.) is a widespread conifer that is the dominant species in woodlands at mid-elevations across a large swath of southwestern North America (West, 1999; Romme et al., 2009). It is an ideal species for the study of genomic variation related to climatic gradients because its current distribution spans 10 degrees of latitude (∼1100 km), elevations ranging from 1500 m to 2500 m (West, 1999), and precipitation regimes from predominantly summer precipitation in southern populations to predominantly winter precipitation among northern populations.

#### 2.1.2 Field sample collection

All trees were sampled in February 2022 at three sites in New Mexico, USA, with each site representing unique points on gradients in latitude and elevation (Fig. 1; Table 1): one in the Jemez Mountains (PO2) and two in the Magdalena Mountains (MG2, MG3). At each site, we selected three mature piñon pines representative of the surrounding stand in size, prioritizing isolated trees to reduce the likelihood of sampling roots from neighboring trees. For each tree sampled, we recorded GPS coordinates, measured diameter and height, and collected an herbarium voucher specimen deposited at the University of New Mexico Herbarium (UNM) (Appendix S1; see Supporting Information with this article). We harvested four tissue types from each adult tree: needles, vegetative buds, phloem, and fine roots (Table 1). Needles and vegetative buds were collected by hand or with scissors. Phloem was sampled by peeling away a bark window and using a razor blade and scalpel to separate phloem tissue from bark and xylem. To collect fine roots, we dug a trench near the base of the tree to locate fine roots, then traced them back to a lateral root where possible to confirm they belonged to the target tree. We also harvested needles from a single wild piñon pine seedling (MG2-RS1-N1; Table 1). Harvested tissues were placed in RNase-free 1.5-mL tubes and immediately frozen on dry ice (PO2) or in liquid nitrogen (MG2, MG3), both of which adequately preserve RNA for field collections (Nagy, 2010; Jordon-Thaden et al., 2015; Yang et al., 2017), then stored at -80° C freezer until further processing.

**Figure 1.**
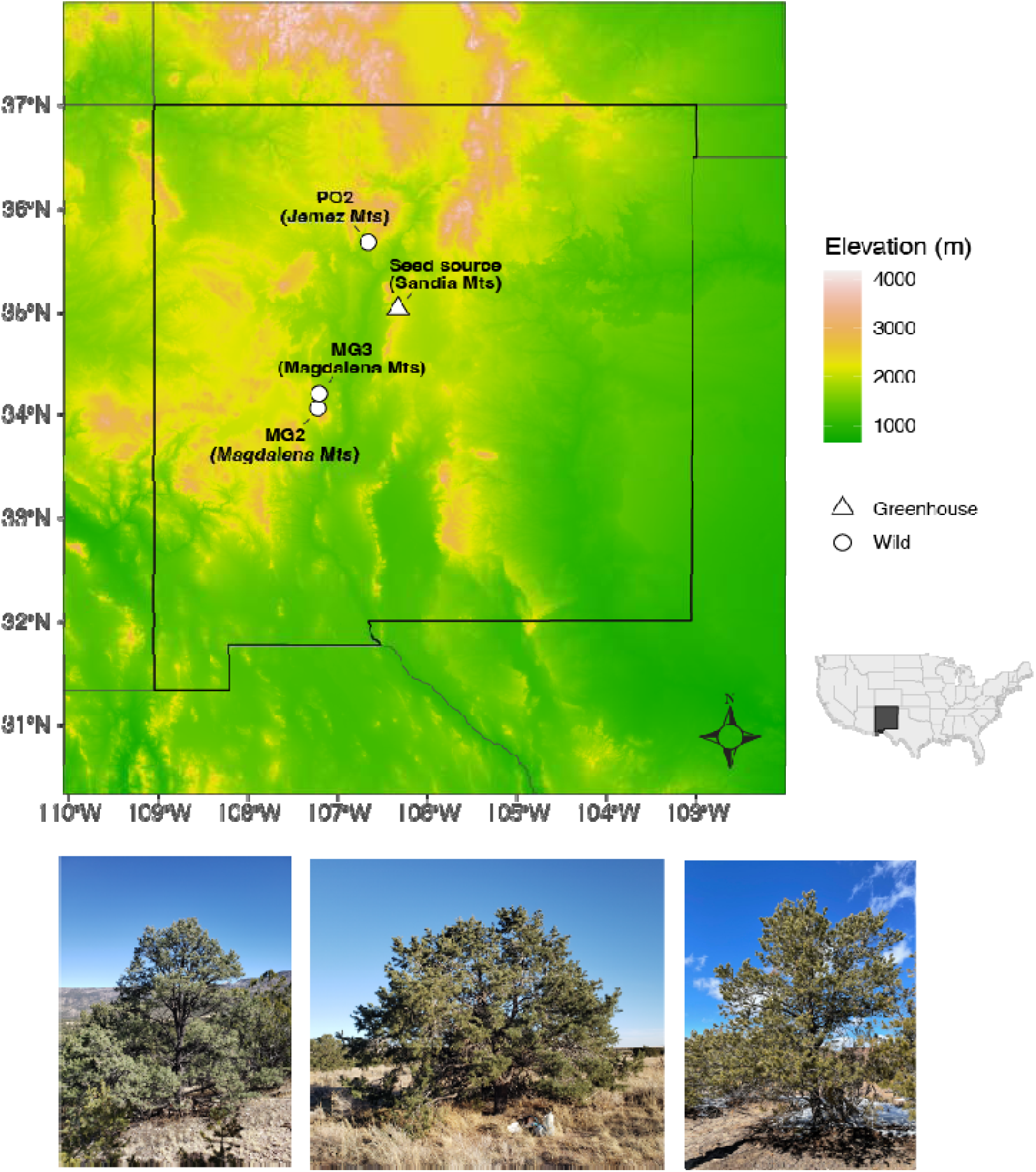
Sampling localities for *Pinus edulis* in New Mexico, USA. Top: elevation map showing the three field sites sampled in February 2022: PO2 in the Jemez Mountains and MG2 and MG3 in the Magdalena Mountains. Seeds collected in the Sandia Mountains in Fall 2020 were used for greenhouse-grown seedlings and megagametophyte tissue. Bottom: representative trees from the three field sites, shown from left to right MG2, MG3, and PO2.

**Table 1.**
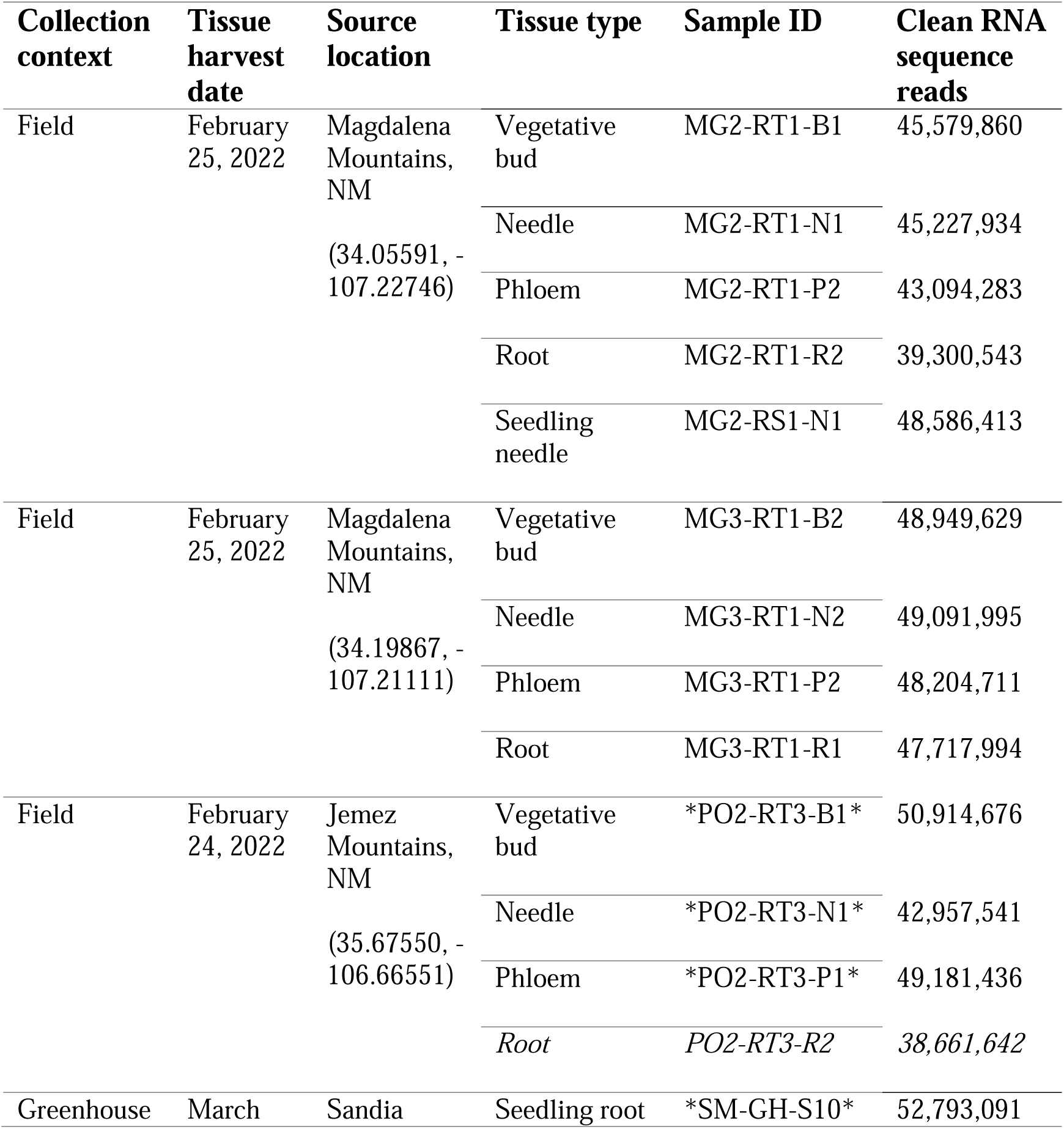
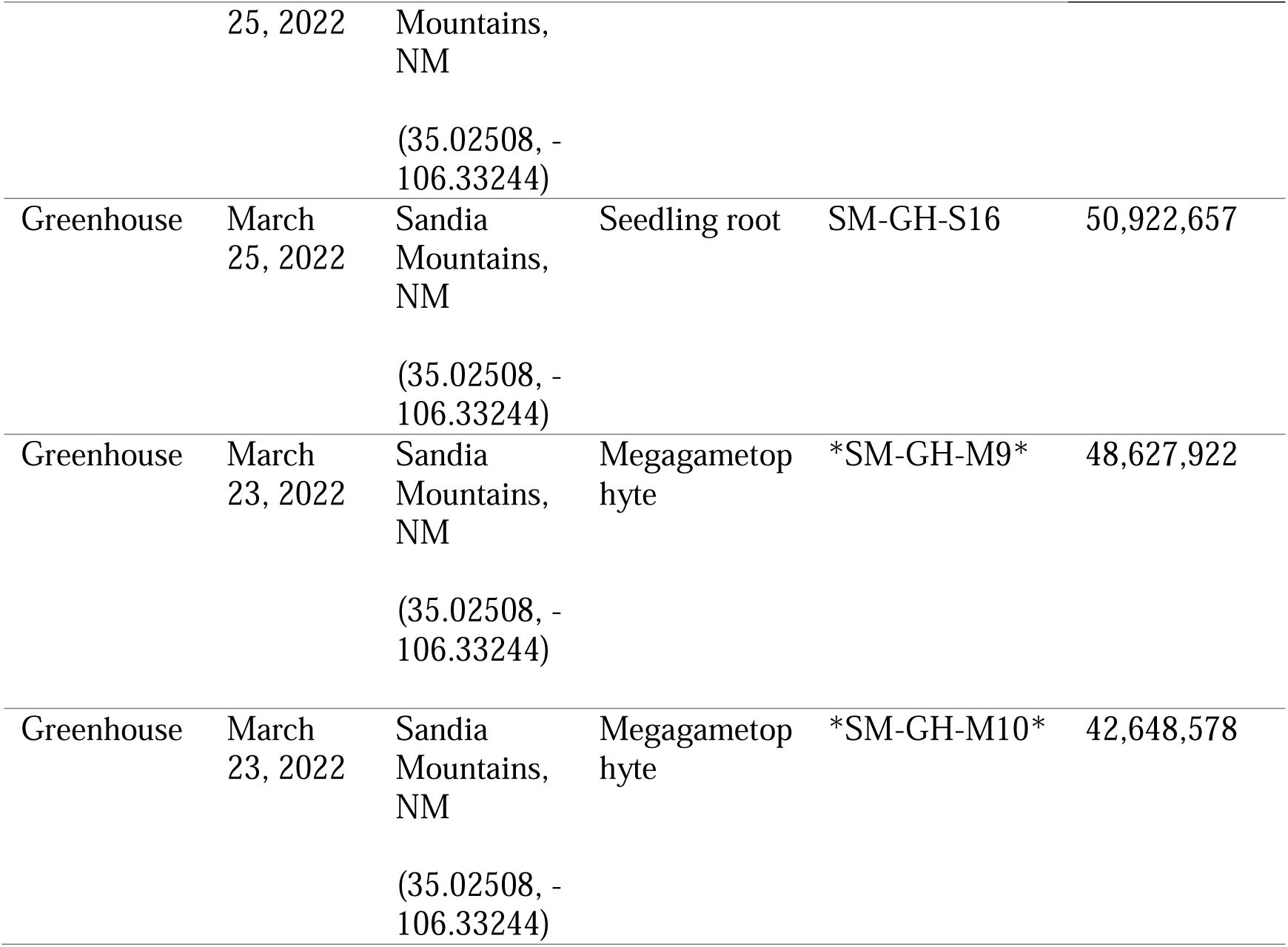
RNA-seq sample information. Collection context, tissue harvest date, source location, tissue type, sample ID, and number of cleaned RNA-seq reads are reported for each library. Location and coordinates for greenhouse samples indicate the seed-source population. RNA-seq libraries represent eight biological sources across multiple tissue types and developmental stages. Asterisks mark the six libraries used to construct the draft reference transcriptome. One library (PO2-RT3-R2, italicized) was excluded because of an outlier expression profile. The remaining 16 libraries were used for downstream expression analyses and probe-target selection.

#### 2.1.3 Greenhouse seedling root sampling

To include genes expressed in early developmental stages, we sampled roots from greenhouse-grown seedlings. We grew seedlings from stored seeds because piñon pines produce seed at irregular intervals, making fresh seeds difficult to obtain in any given year (Chambers et al., 1999; Parmenter et al., 2018). The seeds were collected in September and October 2020 on private property adjacent to the Sandia Mountains, NM, USA, and stored at -20 °C (Table 1). In February 2022, we surface-sterilized 50 seeds in a 10% bleach solution and planted them in an aluminum tray of vermiculite media sealed with cling wrap. The tray was kept on a greenhouse counter at the University of New Mexico and misted twice weekly with sterile water. After one month, we harvested seedling roots, froze them in liquid nitrogen, and stored them at -80 °C until further processing.

#### 2.1.4 Seed megagametophyte sampling for haploid RNA

In March 2022, we obtained haploid megagametophyte tissue from three mature seeds from the collection described above. To induce germination, we placed surface-sterilized seeds on moist Whatman Grade 1 filter paper (Cytiva, Marlborough, MA, USA) in petri dishes. After 24 hours, we dissected the actively growing seeds, using a scalpel and tweezers to remove the diploid nucellar and embryo tissue. The three samples were frozen in liquid nitrogen and stored at -80 °C until further processing.

#### 2.1.5 RNA extractions and sequencing

We pulverized each sample to a fine powder in liquid nitrogen using a mortar and pestle. Total RNA was extracted from approximately 0.1-0.86 g of tissue using the Spectrum Plant Total RNA Kit (Sigma-Aldrich Co., St. Louis, Missouri, USA) following the manufacturer’s protocol with slight modifications to incubation times (e.g., longer tissue digestion times) known to increase RNA quality and yield for conifers (D. O. Vidakovic, personal communication, December 20, 2021). The concentration of RNA was determined by a Qubit 3.0 Fluorometer (ThermoFisher Scientific). The integrity and purity of RNA were assessed with an Agilent 2100 Bioanalyzer (Agilent Technologies, Inc.) equipped with an RNA Nano chip. The RNA quality and yield were validated by the visualization of RNA band integrity by agarose gel electrophoresis.

We selected 17 RNA samples for sequencing based on RNA quality and representation of each tissue type and developmental stage. These included 13 from field collections and four from the seed collection—two greenhouse-grown seedling roots and two megagametophytes (Table 1). Purified RNA from these samples was sent to the Oklahoma Medical Research Foundation Clinical Genomics Center, where RNA-seq libraries were prepared and sequenced on one lane of an Illumina NovaSeq S4 with paired-end 150-bp read output. Following inspection of expression profiles, one library (PO2-RT3-R2) was excluded as an extreme outlier, leaving 16 libraries for downstream analyses.

### 2.2 Reference transcriptome assembly and probe design

All statistical analyses and data visualization were conducted in R v4.4.1 (R Core Team, 2024); bioinformatic analyses were performed using the software and versions specified below.

#### 2.2.1 Read processing and reference transcriptome assembly

To produce a representative set of probes, we used a three-stage process: reference assembly, assembly cleaning, and expression-level filtering for probe design (Fig. 2). First, the quality of paired-end reads was assessed using both FastQC v0.11.9 (Andrews, 2010) and FastP v0.23.2 (Chen et al., 2018). Reports from FastQC were consolidated and visualized together using MultiQC (Ewels et al., 2016). FastP was used to perform quality control for all samples by removing adapter sequences, low quality sequences, and sequences less than 50 base pairs long (option -l).

**Figure 2.**
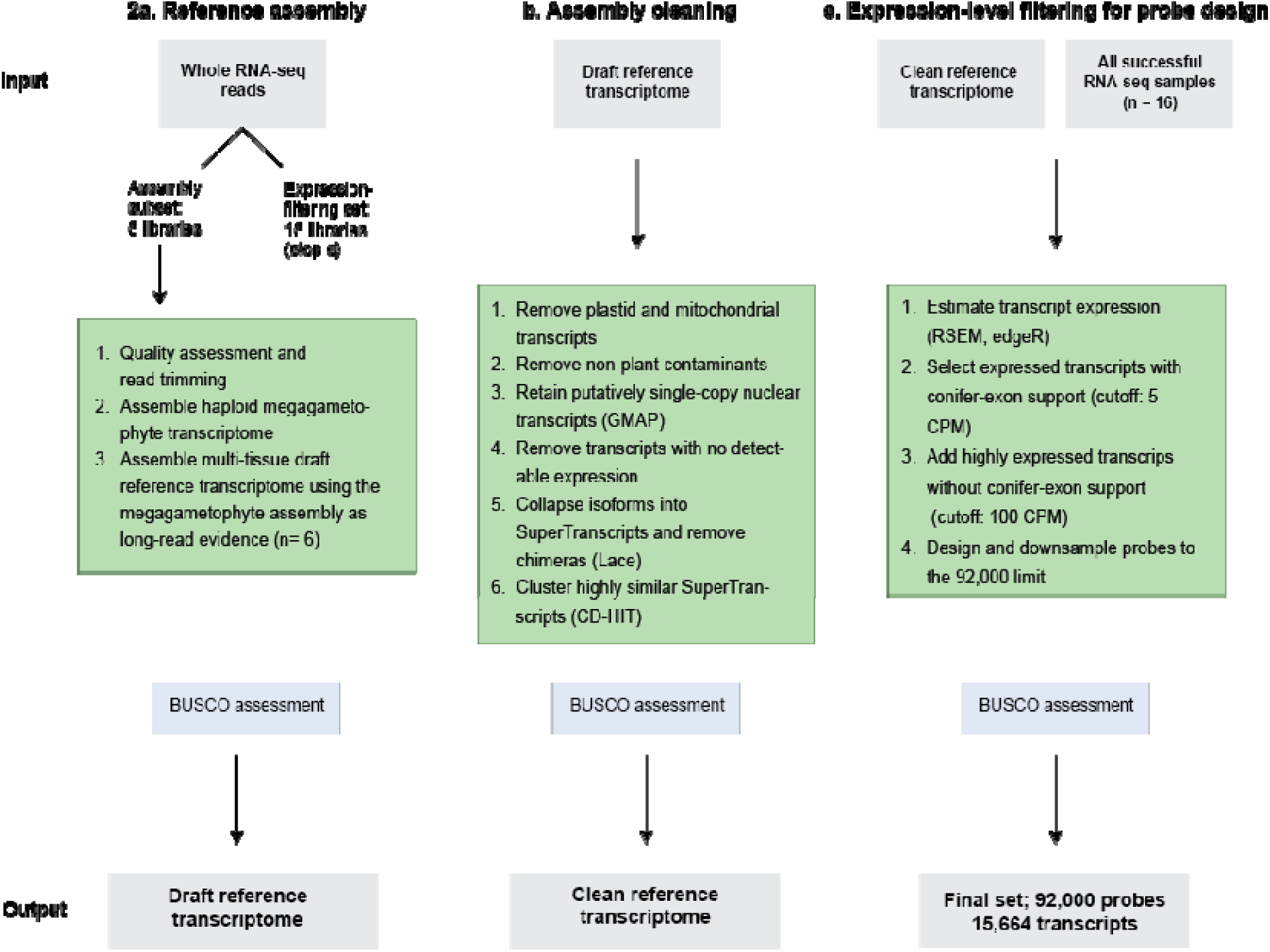
Three-stage workflow from RNA-seq reads to a target-capture set of 92,000 RNA probes targeting 15,664 transcripts for *Pinus edulis*: (a) reference transcriptome assembly, (b) assembly cleaning, and (c) expression-level filtering for probe design. Assembly used a representative subset of libraries spanning focal tissue types. Reads from all 16 retained libraries were then mapped to the clean reference transcriptome for expression-level target selection. Transcripts with conifer-exon matches were retained at ≥5 CPM, whereas those without were required to reach ≥100 CPM before probe design and downsampling to the 92,000-probe limit. Assembly and target sets were assessed with BUSCO.

We used a pseudo-guided transcriptome assembly approach, following Ojeda et al. (2019), to build a draft reference transcriptome. This approach uses a transcriptome assembled from haploid tissue as long-read evidence to guide an assembly incorporating multiple tissue types. We used this approach because haploid-derived sequences are expected to improve assembly reliability due to the lack of heterozygosity, while still incorporating transcripts from all focal tissue types. We first assembled a *de novo* transcriptome using the reads from *Pinus edulis* haploid megagametophyte tissue (two individuals; Table 1) using Trinity v2.13.2 (Grabherr et al., 2011). We then assembled a draft reference transcriptome in a single Trinity run using reads from adult needle, bud, and phloem tissues collected from one field individual and greenhouse-grown seedling root tissues (Table 1). For this second assembly, we used the --long_reads option to incorporate the megagametophyte transcriptome assembly as if it were long-read evidence (Ojeda et al., 2019). We assessed draft transcriptome completeness using Benchmarking Universal Single-Copy Orthologs (BUSCO) v5.1.3 (Simão et al., 2015) with the plant database from OrthoDB v10 (Manni et al., 2021), which assesses completeness based on the presence of universal single-copy orthologs. We calculated N50 for all contigs using QUAST v5.0.2 (Gurevich et al., 2013).

#### 2.2.2 Assembly cleaning

Because our aim was to design probes targeting putatively single-copy regions of the *P. edulis* nuclear genome, we cleaned the draft reference transcriptome in several stages. These filters were used to remove: (1) plastid and mitochondrial sequences, (2) non-plant contaminants sequenced alongside *P .edulis*, (3) transcripts likely derived from multi-copy nuclear gene regions, (4) transcripts without detectable expression, and (5) transcript-level redundancy.

We first identified organellar sequences using BLASTN v2.13.0 (Altschul et al., 1990) against the *Pinus lambertiana* chloroplast genome (NCBI GenBank accession NC_011156) and the *Pinus taeda* mitochondrial genome (NCBI GenBank accession NC_039746.1). Transcripts that matched either genome with an e-value cutoff of 1e-10 were removed from the draft reference transcriptome.

We screened the remaining transcripts for non-plant contaminants using DIAMOND BLASTX v2.0.15 (Buchfink et al., 2021) against the NCBI non-redundant protein database. We removed transcripts that mapped only to non-plant proteins with an e-value cutoff of 1e-10. Transcripts that mapped to both plant and non-plant sequences with the same e-value cutoff were retained. We then aligned the remaining transcripts to the *P. lambertiana* genome (NCBI GenBank accession GCA_001447015.2) using GMAP v2021.08.25 (Wu and Watanabe, 2005) and retained only transcripts with a single genomic hit to prioritize putatively single-copy nuclear regions.

We next excluded transcripts without detectable expression across any sample, as estimated using RSEM v1.3.3 (Li and Dewey, 2011) and edgeR v3.38.4 (Robinson et al., 2010). This presence–absence filter removed unsupported transcripts and was distinct from the cpm threshold filtering used for probe-target selection in Section 2.2.3.

To reduce redundancy arising from isoforms, allelic variants, fragmented transcripts, and overlapping contigs, we generated SuperTranscripts using Lace v1.14.1 (Davidson et al., 2017), following Ojeda et al. (2019). SuperTranscripts combine multiple isoforms into a single non-redundant contig representing the expressed sequence of a gene. We removed potentially chimeric SuperTranscripts, defined here as those with multiple BLAST hits within themselves, and clustered the remaining sequences using CD-HIT v4.8.1 (Li and Godzik, 2006; Fu et al., 2012) to collapse highly similar sequences.

Several filtering steps were performed with the subseq module in seqtk v1.3-r106 (Li, 2012), which allowed us to subset FASTA files to loci retained at successive stages of workflow (Fig. 2, Tables 2, 3). We assessed the reference transcriptome using BUSCO v5.1.3 (Simão et al., 2015) before, during, and after cleaning to evaluate how each filtering step affected conserved gene content and duplication (Table 2).

**Table 2.** Effect of sequential assembly-cleaning steps on reference transcriptome completeness, duplication, and assembly size. BUSCO results are reported as the percentage of complete orthologs (C), including single-copy (S) and duplicated (D) orthologs. Each row summarizes the assembly retained after the indicated filtering or clustering step. The final row represents the clean reference transcriptome.

| **Description** | **BUSCO (C: S, D)** | **Transcripts** | **Base pairs** | **Mean sequence length (bp)** |
| --- | --- | --- | --- | --- |
| Haploid-guided assembly after decontamination | 91.4%: 20.0%, 71.4% | 605,272 | 367,349,431 | 606.92 |
| Transcripts with a single hit to reference | 84.4%: 20.1%, 64.3% | 260,174 | 205,576,152 | 790.15 |
| Remove sequences without detectable expression | 84.4%: 20.2%, 64.2% | 241,018 | 199,568,514 | 828.02 |
| Lace supertranscripts (chimeras removed) | 76.1%: 73.4%, 2.7% | 152,659 | 84,307,600 | 552.26 |
| *CDHIT clustered clean (final reference transcriptome)* | *76.0%: 73.4%, 2.6%* | *148,910* | *83,106,474* | *558.10* |

**Table 3.** Effects of alternative combinations of database-match criteria and expression thresholds on the transcript subset considered for probe design. BUSCO completeness (C), relative proportions of single-copy (S) and duplicated (D) loci, and summary statistics are shown for each subset. Each filtering strategy combined *Pinus*- or conifer-matched transcripts with highly expressed nonmatching transcripts (≥100 cpm in at least one sample). *Pinus* matches included transcripts corresponding to reference exons from *Pinus lambertiana* or *P. taeda*, whereas conifer matches additionally included *Picea abies* and *P. glauca*.

| Description | BUSCO [C: S, D] | Transcripts | Base pairs | Mean sequence length (bp) | Probe density (target bp/probe) |
| --- | --- | --- | --- | --- | --- |
| Reference transcriptome | 76.0%: 73.4%, 2.6% | 148,910 | 83,106,474 | 558.10 | 903.33 |
| *Pinus* matches, no cpm cutoff + nonmatching transcripts $\geq 100$ cpm | 74.6%: 72.0%, 2.6% | 54,711 | 48,771,891 | 891.45 | 530.13 |
| *Pinus* matches $\geq 5$ cpm + nonmatching transcripts $\geq 100$ cpm | 74.2%: 71.6%, 2.6% | 17,555 | 33,091,381 | 1885.01 | 359.69 |
| Conifer matches, no cpm cutoff + nonmatching transcripts $\geq 100$ cpm | 76.0%: 73.4%, 2.6% | 57,294 | 50,231,775 | 876.74 | 546.00 |
| Conifer matches $\geq 5$ cpm + nonmatching transcripts $\geq 100$ cpm | 75.5%: 72.9%, 2.6% | 17,913 | 33,500,238 | 1870.16 | 364.13 |
| *Final probe design* | *75.2%: 72.7%, 2.5%* | *15,664* | *31,392,557* | *2004.12* | *341.22* |

#### 2.2.3 Expression level filtering and probe design

We used expression-level filtering to reduce the cleaned reference transcriptome to a feasible target size for probe design while prioritizing expressed and functionally supported transcripts (Fig. 2; Table 3). Transcript abundance was estimated by aligning reads from all 16 retained libraries to the cleaned reference transcriptome using RSEM v1.3.3 (Li and Dewey, 2011) and edgeR v3.38.4 (Robinson et al., 2010), parallelized with GNU Parallel v20190222 (Tange, 2018).

Expression-level filtering was based on counts-per-million-reads (cpm). At each evaluated threshold, a transcript was retained when at least one sample had expression above the threshold, allowing transcripts expressed strongly in a single tissue type, life stage, or individual to be retained. We evaluated multiple combinations of cpm thresholds and functional support from a conifer exon database, as summarized in Table 3.

For the final filtering strategy, we retained transcripts that had both a DIAMOND BLASTX v2.0.15 (Buchfink et al., 2021) match to the conifer exon database and expression of at least 5 cpm in one or more *P. edulis* samples from the present study. The conifer species included in the exon database were *Picea abies* (Nystedt et al., 2013), *Picea glauca* (Warren et al., 2015), *Pinus lambertiana* (Stevens et al., 2016) and *Pinus taeda* (Zimin et al., 2017). Second, to retain highly expressed transcripts that were not represented in the conifer exon database, we also included transcripts expressed at ≥100 cpm in at least one sample. Because this filtered set exceeded the predetermined probe limit, probes were downsampled across eligible transcripts during final probe design performed by Rapid Genomics (Gainesville, Florida, USA; now part of LGC Biosearch Technologies) using a design limit of 92,000 120-bp probes.

### 2.3 Reference transcriptome and probe-set assessment

#### 2.3.1 Tissue-specific expression

To assess the validity of the reference transcriptome and variation in expression profiles among samples and tissue types, we used multidimensional scaling (MDS) in edgeR (Robinson et al., 2010) using the RSEM expression estimates from all 16 samples that passed sequencing quality control (Fig. 3).

**Figure 3.**
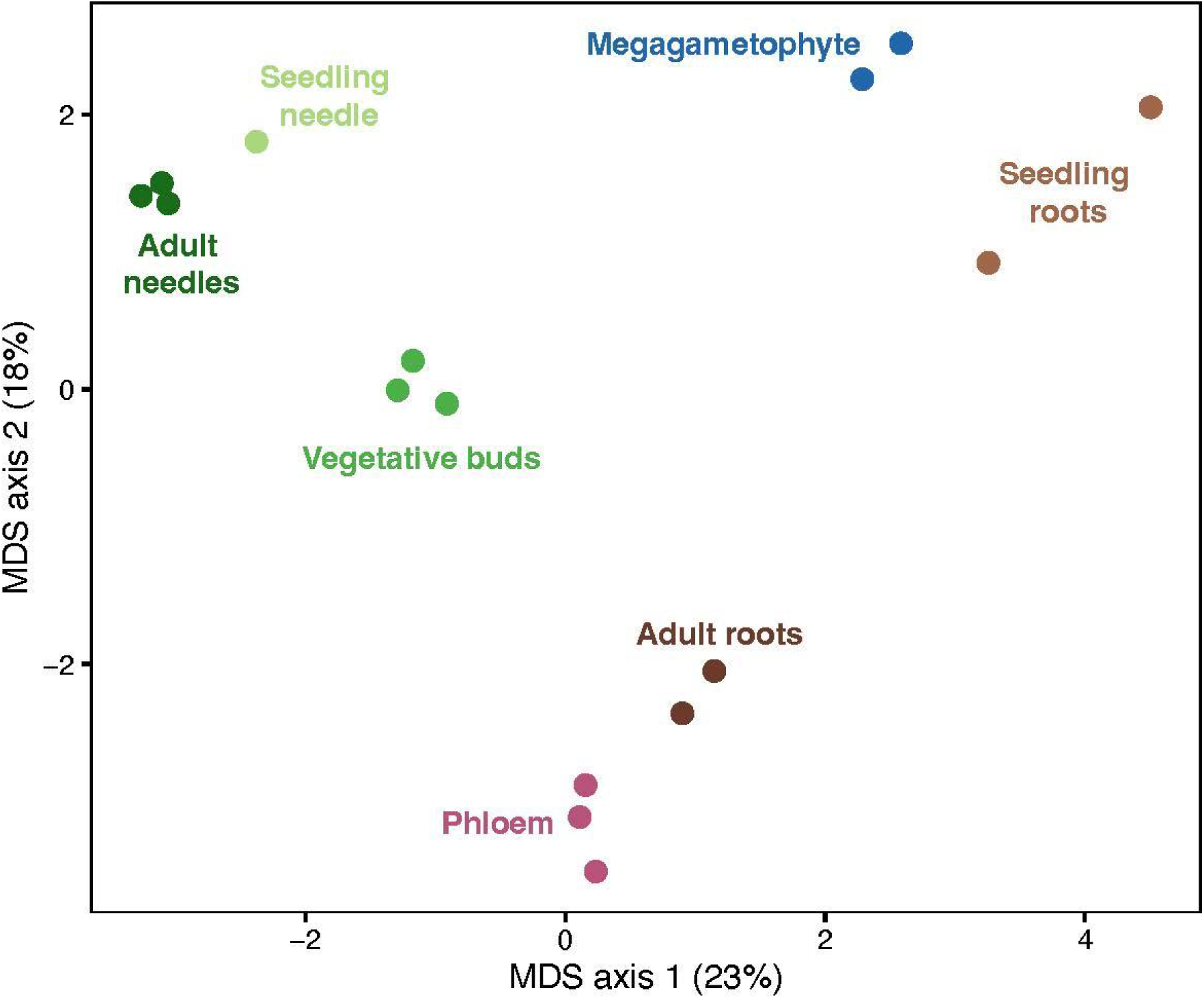
Multidimensional scaling of expression profiles from the 16 retained RNA-seq libraries. Colors indicate tissue-stage groups. Seedling roots and megagametophytes were greenhouse-grown; all other tissue types were field collected. Distances represent leading log-fold-change differences among libraries. The first and second dimensions accounted for 23% and 18% of the variation in these distances, respectively. Samples group primarily by tissue, with photosynthetic aboveground tissues (adult needles, seedling needles, and vegetative buds) grouping together.

#### 2.3.2 Contributions of multiple tissue types to probe set

To quantify the contribution of each tissue to the final probe set, we evaluated seven tissue-stage groups: adult needles, seedling needles, adult phloem, adult vegetative buds, adult roots, seedling roots, and megagametophytes. For each group, we identified probe-set transcripts expressed above 5 cpm in that group but not in any other group. We interpreted these as the unique contribution of each tissue-stage group and expressed as a percentage of the 15,664 targeted transcripts.

To address overlap among functionally similar tissues, we repeated this analysis using broader functional categories. Photosynthetic tissues included adult needles, seedling needles, and vegetative buds, whereas root tissues included adult and seedling roots. For each category, we counted transcripts expressed exclusively within that category. We calculated within-category overlap as the difference between this value and the summed unique contributions of its constituent tissue-stage groups.

As a robustness analysis, we estimated the average marginal contribution of each tissue-stage group across all possible combinations of the seven groups. For each combination, we counted the number of probe-set transcripts expressed above 5 cpm in at least one included group. We then fit a linear model of this count against the binary inclusion of each tissue-stage group. The resulting coefficients estimated the mean number of transcripts added by each tissue-stage group across all combinations.

We visualized the shared and unique expression among four focal tissue types using Venn diagrams made with ggvenn v0.1.10 (Yan, 2021). These included the three highest-ranked tissue types from the contribution analyses—seedling roots, phloem, and adult needles—as well as megagametophytes. In contrast to the contribution analyses, the Venn diagrams included all transcripts in the full reference transcriptome expressed above 5 cpm in at least one sample of each tissue type.

#### 2.3.3 Functional annotation of highly expressed tissue-stage transcripts

To evaluate whether tissue-specific expression patterns reflected plausible biological differences among the tissues and life stages, we identified highly enriched transcripts within each of the seven tissue-stage groups. For each transcript, we calculated mean counts per million reads (cpm) within each tissue-stage group and identified the group with the highest mean expression. Transcripts were considered highly enriched when their highest mean expression exceeded 1,000 cpm and was at least 10-fold greater than that of the next-highest group.

We retained up to five highly enriched transcripts per tissue-stage group for functional annotation. Sequences were extracted from the reference transcriptome using the subseq module in seqtk v1.3 (Li, 2012), and queried with BLASTX (Altschul et al., 1990) against the NCBI RefSeq protein database restricted to Viridiplantae (taxid 33090). We retained the highest-scoring match for each transcript and considered annotations confident when they matched a named, non-hypothetical protein with ≥ 50% amino-acid identity. Other matches were treated as tentative.

#### 2.3.4 Detecting paralogs

To identify transcripts potentially affected by paralogy, we used reads from the haploid megagametophytes to screen for unexpected heterozygous SNP calls. Because true heterozygosity is not expected in haploid tissue, such calls may indicate collapsed paralogs, assembly artifacts, or problematic read mapping.

We called variants after aligning megagametophyte reads to both the unfiltered reference transcriptome and the filtered reference transcriptome used for probe design. We used a standard pipeline, first aligning reads with BWA-mem v0.7.17 (Li and Durbin, 2009), then removing unmapped reads and converting the output to BAM files with samtools v1.11 (Li et al., 2009), marking duplicates with Genome Analysis Toolkit v4.1.9 (McKenna et al., 2010), and finally indexing the BAM files with samtools. These BAM files were then used as input for bcftools v1.12 (Li, 2011). Heterozygous SNP calls were counted for each transcript, and transcripts with heterozygous calls at more than 1% of evaluated sites in both megagametophyte samples were flagged as potentially affected by paralogy.

## 3. RESULTS

### 3.1 Sequencing

Across the 17 RNA-seq libraries, sequencing yield ranged from 38.7 to 52.8 million paired-end reads, and base-quality distributions consistently peaked at a Phred score of 36. Sequencing depth was relatively consistent across libraries and sufficient for reference transcriptome assembly and downstream expression analyses (Table 1). We excluded one adult-root library from the Jemez Mountains (PO2-RT3-R2) from downstream expression analyses because it had an extreme expression profile: 80.3% of its estimated transcript abundance was concentrated in three transcripts, compared with less than 20% in the three most abundant transcripts of every other library. The remaining 16 libraries were retained for transcriptome assessment, expression filtering, and tissue-expression analyses.

### 3.2 Reference transcriptome assembly and probe design

The haploid-guided draft reference transcriptome recovered 91.4% of single-copy plant genes but, as expected for a *de novo* conifer transcriptome, contained substantial duplication (71.4% duplicated BUSCOs; Table 2). Because our goal was to design probes targeting putatively single-copy nuclear regions, we applied sequential filters to remove organellar and contaminant sequences, exclude putatively multicopy loci, and reduce transcript-level redundancy (Fig. 2; Table 2). These steps reduced duplicated BUSCOs to 2.6% while retaining 76.0% complete BUSCOs, producing a clean reference transcriptome of 148,910 transcripts.

The cleaned transcriptome (148,910 transcripts) was still far too large for our probe budget of 92,000 probes (120 bp each, capturing ∼150 bp per probe), so we used expression-based filtering to prioritize informative targets (Fig. 2; Table 3). We retained transcripts with both a conifer-database match and expression above 5 cpm, as well as transcripts without a database match when expression exceeded 100 cpm. This filtering reduced the target set approximately eightfold to 17,913 transcripts spanning 33.5 Mbp while retaining 75.5% complete BUSCOs. This set still exceeded our probe budget, so during synthesis we evenly downsampled probes from the highest-coverage transcripts. The 2249 transcripts that received no probes were shorter than those retained (median 767 versus 1819 bp), suggesting that their exclusion was associated with insufficient target length. The final probe set targeted 15,664 transcripts (31.4 Mbp; ∼0.1% of the genome), with each transcript represented by one or more of the 92,000 probes. BUSCO completeness remained stable at 75.2% in the final set, indicating that target reduction preserved most conserved gene content.

### 3.3 Expression profiles across tissues and developmental stages

The first two MDS dimensions accounted for 23% and 18% of the variation in leading log-fold-change distances among samples, respectively (41% combined; Fig. 3). Expression profiles separated primarily by tissue type and developmental stage. Adult and seedling needles clustered near vegetative buds, indicating similar expression profiles among photosynthetic tissues. Adult roots and phloem also clustered together, whereas seedling roots and megagametophytes formed a separate grouping. Replicate samples from the same tissue-stage group generally occurred near one another, supporting the consistency of the expression profiles used to construct and evaluate the reference transcriptome.

### 3.4 Contributions of multiple tissue types

Every tissue type contributed transcripts that were not expressed above 5 cpm in any other tissue type (Fig. 4A). Seedling roots made the largest unique contribution, with 1187 transcripts representing 7.6% of the 15,664 probe-set transcripts. They were followed by phloem (378; 2.4%), adult needles (279; 1.8%), vegetative buds (250; 1.6%), megagametophytes (231; 1.5%), adult roots (161; 1.0%), and seedling needle (26; 0.2%). In total, 2512 probe-set transcripts, or 16.0%, of the final probe set, were expressed above the threshold in only one tissue-stage group.

**Figure 4.**
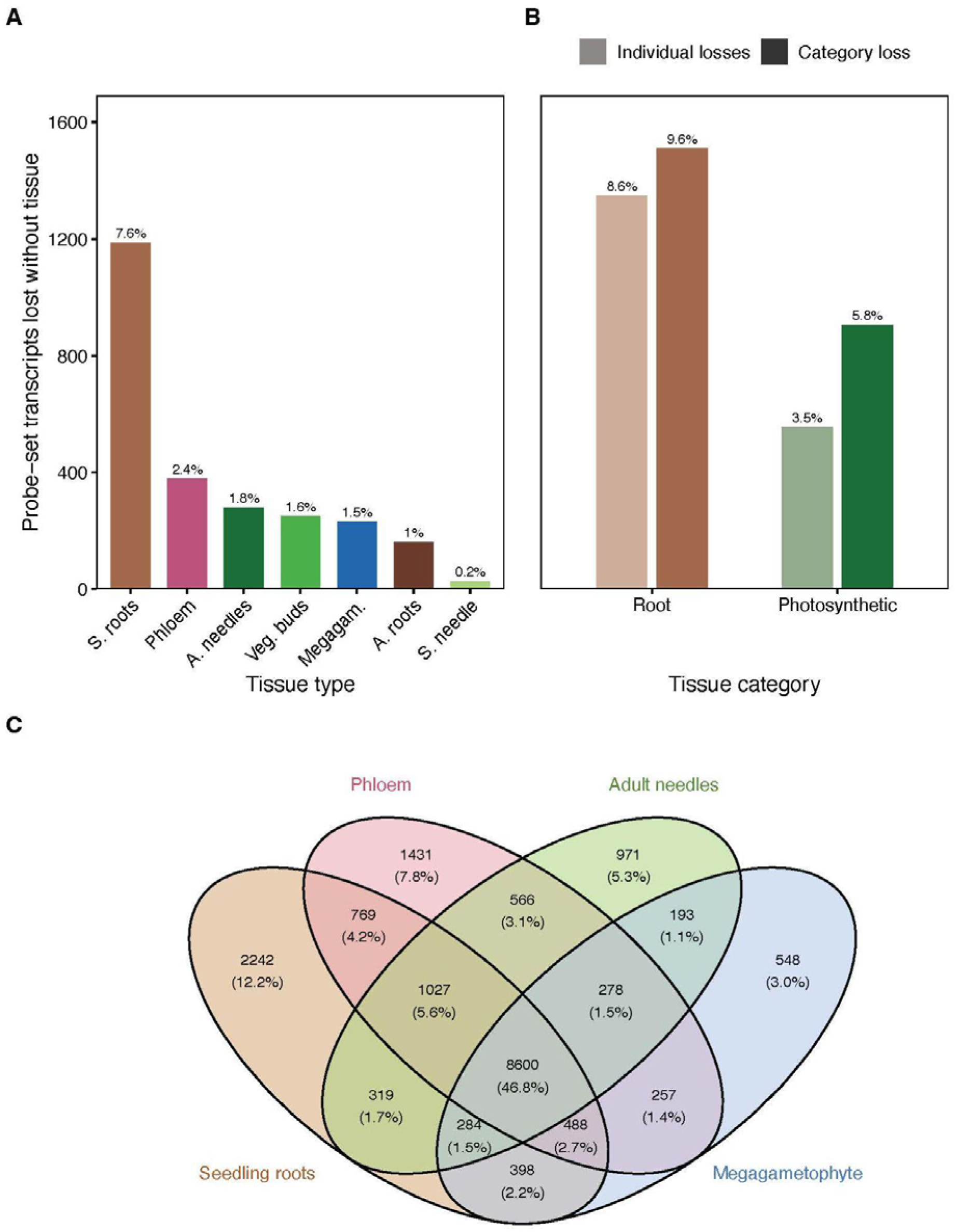
Quantifying the contribution of each tissue to the probe set. (A) Probe-set transcripts lost if each tissue were excluded, shown as a count and as a percentage of the 15,664 transcripts in the final probe set. “S” denotes seedling and “A” denotes adult. (B) Transcripts lost if an entire tissue category were excluded (solid bars), compared with the sum of the individual tissue losses within that category (faded bars); the gap reflects transcripts shared among tissues within a category. Only categories sampled at more than one tissue group are shown: Roots (adult and seedling) and Photosynthetic (adult needles, vegetative buds, and seedling needles). (C) Venn diagram of unique and shared transcripts expressed (cpm >5) across four focal tissue types: the three highest-ranked in (A) plus the megagametophyte. Unlike (A) and (B), which evaluate transcripts in the final probe-set, (C) evaluates the broader set of transcripts expressed in at least one of the four focal tissue types (18,371).

Grouping tissue types into broader functional categories revealed substantial redundancy within one category but not others (Fig. 4B). Photosynthetic tissue (adult needles, vegetative buds, seedling needle) collectively contributed 5.8% of the probe set, compared with 3.5% when their individual unique contributions were summed. This difference represents transcripts shared among multiple photosynthetic tissue-stage groups but not expressed above the threshold in other tissue categories. Root tissues showed little such redundancy (9.6% as a category vs. 8.6% summed), indicating that adult and seedling roots have less within-category overlap and more unique contributions per tissue type. The average marginal-contribution analysis across all combinations of tissue-stage groups supported the same overall pattern. Seedling roots had the largest estimated average contribution (1855 transcripts; 11.9% of the probe-set union), followed by phloem (987; 6.3%), whereas seedling needles had the smallest contribution (413; 2.6%). Vegetative buds ranked above adult needles by average marginal contribution but below adult needles by unique contribution, consistent with the greater expression overlap among photosynthetic tissues shown in Figure 4B. In the broader set of 18,371 transcripts expressed across the four focal tissues, 8600 (46.8%) were shared among all four, whereas seedling roots had the largest tissue-specific set (2242 transcripts; 12.2%; Fig. 4C).

We also evaluated how much of the final probe set would have been represented by needle tissue alone. Of the 15,664 probe-set transcripts, 11,365 (72.6%) were expressed above 5 cpm in at least one adult or seedling needle sample. The remaining 4299 transcripts (27.4%) were detected only in non-needle tissues and would not have been represented using the same expression criterion in a needle-only design. Restricting the comparison to adult needles, as is typical in conifer phylogenomic studies, would have excluded a larger fraction, 4675 transcripts or 29.8%, of the final probe set.

### 3.5 Functional annotation of highly enriched tissue-stage transcripts

Of the 29 transcripts selected for annotation, 17 had confident matches to named proteins and are presented in Table 4. The clearest functional signals came from those that were most transcriptionally distinct (Figs. 3, 4). Megagametophyte-enriched transcripts included genes associated with seed-metabolism, germination, and desiccation. These included glyoxylate-cycle enzymes isocitrate lyase and malate synthase, which had 85.7% and 85.6% identity and contribute to the mobilization of stored lipids during germination (Eastmond and Graham, 2001; Graham, 2008). We also identified an 11-kDa late embryogenesis abundant (LEA) protein associated with seed desiccation tolerance (Hernández-Sánchez et al., 2022). Phloem-enriched transcripts similarly included defense- and stress-associated genes, including the antimicrobial peptide defensin (Ec-AMP-D1-like), NAD kinase 3 and a metallothionein-like protein associated with redox regulation and oxidative stress (Hassinen et al., 2011; Tam et al., 2015; Li et al., 2018).

**Table 4.**
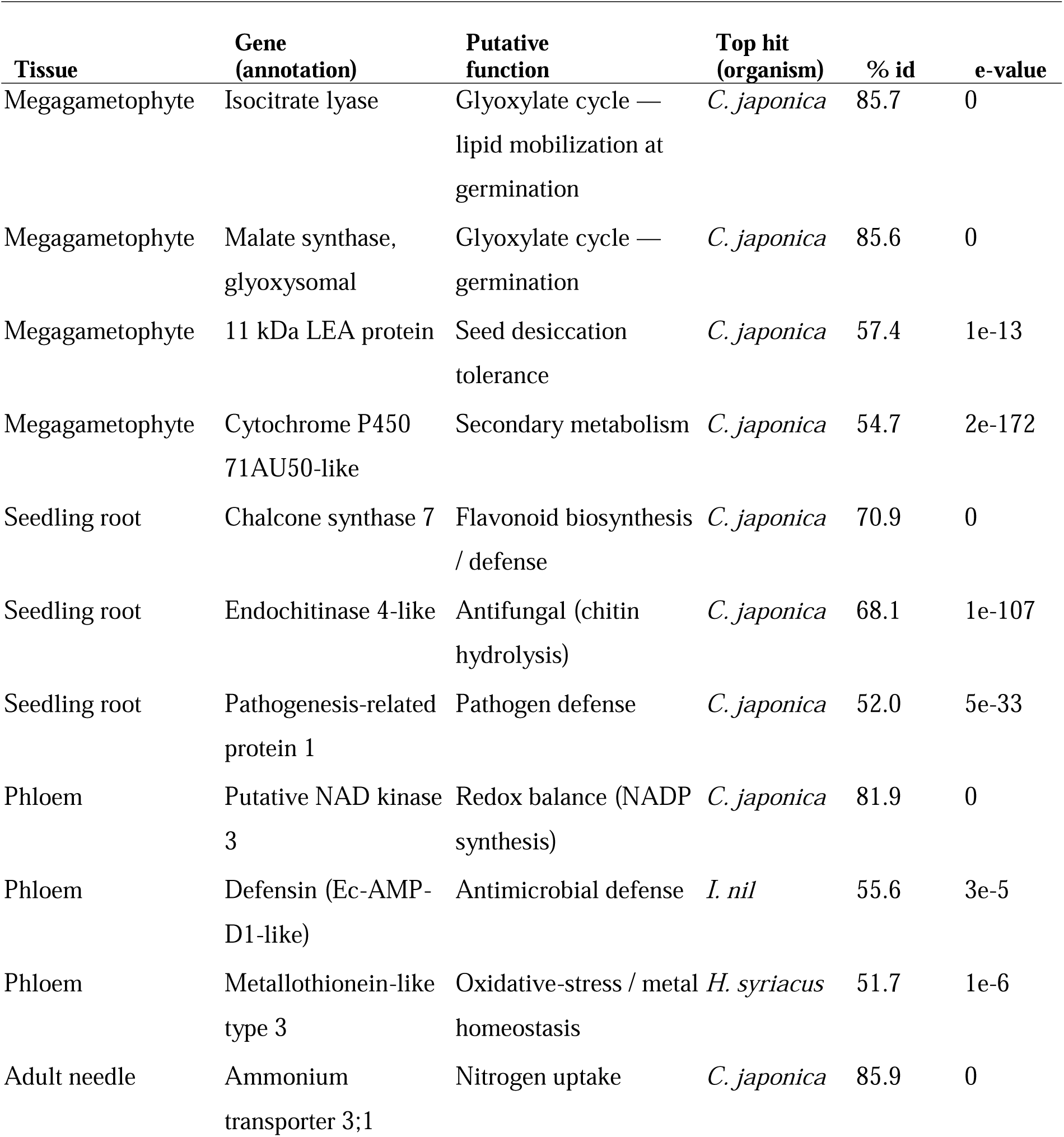
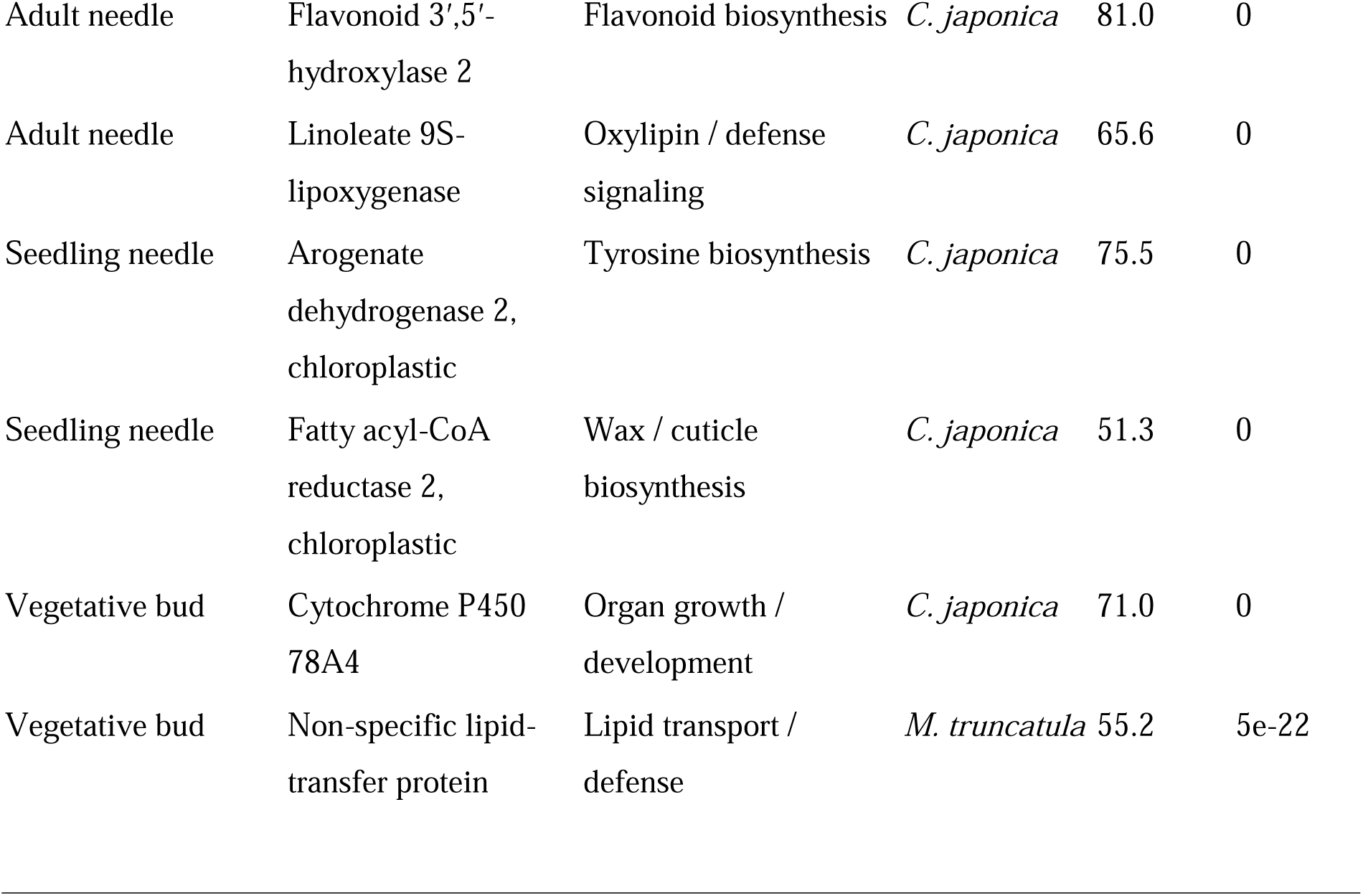
Functional annotation of tissue-specific marker transcripts. Markers were highly expressed (mean >1,000 CPM) and ≥10-fold tissue specific and were annotated by BLASTX against RefSeq Viridiplantae (best hit shown). Only confident annotations (named proteins, ≥50% identity) are listed. Best-hit organisms include: *Cryptomeria japonica, Ipomoea nil, Hibiscus syriacus, and Medicago truncatula*.

Only one transcript met the enrichment criteria for adult roots, and its best match was an uncharacterized protein. However, seedling-root-enriched transcripts included three proteins annotated with defense-related functions: pathogenesis-related protein 1, chalcone synthase, and an endochitinase. Transcripts selected from adult needles, seedling needles, and vegetative buds generally had broader metabolic annotations and fewer clearly tissue-associated functions. This pattern was consistent with the greater expression overlap among photosynthetic tissues observed in the MDS and tissue-category analyses (Figs. 3, 4B).

Most best-scoring matches were to proteins from *Cryptomeria japonica* (Table 4). Several transcripts matched only uncharacterized or hypothetical proteins, consistent with the limited functional annotation currently available for many conifer proteins.

### 3.6 Screening for paralog-associated heterozygosity

The haploid-tissue screen flagged 1590 transcripts as potentially affected by collapsed paralogs, assembly artifacts, or ambiguous read mapping. These represented 8.88% of the 17,913 candidate transcripts retained before final probe downsampling (Table 3) and had heterozygous calls at more than 1% of evaluated sites in both megagametophyte samples. Transcripts without sufficient expression in the megagametophytes could not be assessed using this approach.

## 4. DISCUSSION

Genomic research on conifers has been hindered by long generation times, large genomes, and limited funding (Neale and Kremer, 2011). In the absence of a reference genome for piñon pine, we assembled a multi-tissue reference transcriptome that retained roughly three-quarters of conserved plant genes while representing less than 0.3% of its approximately 30-Gb genome. From this resource, we designed an expression-filtered exon-capture probe set targeting only approximately 0.1% of the genome. Assembling a chromosome-scale genome for such a large, repeat-rich genome remains costly and technically demanding, but is not a prerequisite for targeted capture. Probe design requires only a defined fraction of the genome, and a multi-tissue transcriptome directly recovers expressed sequences that can serve as biologically relevant targets. Although the individual methods used here are established, we integrated them for a goal distinct from that of most existing capture tools. Rather than targeting the conserved, low-copy orthologs favored by phylogenomic bait sets (Weitemier et al., 2014; Khan et al., 2024), our multi-tissue, expression-guided design prioritized genes expressed across organs and life stages (Fig. 2). The resulting probe set is therefore oriented toward population, landscape, and conservation genomics, where variation in expressed genes may help explain ecological performance and responses to environmental change.

The central question motivating this study was how much multi-tissue sampling contributes to a probe set and what would be lost through a conventional single-tissue approach. We found the cost of narrow tissue sampling to be substantial (Fig. 4). Of the 15,664 probe-set transcripts, only 72.6% were expressed in needle tissue, whereas 27.4% were expressed only in non-needle tissues and would not have been recovered using the same expression criteria in a needle-only design. The functional annotations illustrate the types of biological information that broader tissue sampling can recover. Megagametophytes contributed transcripts associated with seed desiccation tolerance and germination, including a late embryogenesis abundant protein and glyoxylate-cycle enzymes (Shih et al., 2008; Hernández-Sánchez et al., 2022). Roots and phloem likewise contributed defense- and stress-associated transcripts, including an antimicrobial defensin in phloem (Picart et al., 2012), highlighting functions that would be poorly represented by foliage alone. Capturing variation in these genes could be especially valuable for landscape-genomic studies of piñon pines, which occur across substantial climatic, edaphic, and biotic variation (Gehring et al., 2017; Whipple et al., 2019; Buck et al., 2023). By contrast, the greater overlap among photosynthetic tissues suggests that expanding sampling across distinct organs and life stages may recover more functionally diverse targets than adding closely related foliage tissues (Figs. 3, 4). A systematic functional analysis (e.g., gene-ontology enrichment per tissue) would formalize these patterns and is a natural next step as conifer annotation improves (Neale and Kremer, 2011).

Beyond probe design, the transcriptomes and RNA-seq data provide an opportunity to compare tissue-level expression patterns across conifer species. Expression profiles in *P. edulis* grouped primarily by tissue type, supporting the biological coherence of the libraries and revealing broad similarities among functionally related tissues. Such comparisons can help distinguish expression patterns that are conserved across conifers from those that may be species- or environment-specific, which is important when transferring candidate genes and genomic tools among closely related pinyon species. For example, needles and vegetative buds clustered closely in *P. edulis* (Fig. 3), whereas Cervantes et al. (2021) reported greater separation between these tissues in *P. sylvestris* using a similar assembly framework (Ojeda et al., 2019). These differences may reflect species-specific expression, but they could also arise from differences in sampling season, developmental stage, or growing conditions. Broader comparative analyses across conifer species and environments will be needed to determine which tissue-expression patterns are general and which are context dependent.

We designed the probe set to support population- and landscape-genomic studies and the discovery of variants associated with ecological performance, reproduction, climate resilience, and other adaptive traits (Suren et al., 2016; Syring et al., 2016; Yeaman et al., 2016). Haploid megagametophyte tissue strengthened this resource in two ways: it improved transcriptome assembly (Ojeda et al., 2019) and provided an independent screen for paralog-associated heterozygosity. Because true heterozygosity is not expected in haploid tissue, transcripts with heterozygous calls in both megagametophyte samples can be flagged as potentially affected by collapsed paralogs, assembly artifacts, or ambiguous read mapping. Although alignment to the related *P. lambertiana* genome helped prioritize putatively single-copy regions, 8.88% of candidate transcripts were still flagged by the haploid screen. Transcripts with adequate megagametophyte coverage and no unexpected heterozygosity can therefore serve as a higher-confidence whitelist, whereas flagged transcripts can be excluded or treated cautiously in downstream exome-capture analyses.

Building this resource required deliberate compromises while preserving its utility for downstream genomic applications (Rellstab et al., 2015; Suren et al., 2016; Isabel et al., 2020). We sampled multiple tissues and life stages, assembled the transcriptome from a representative subset of libraries, and used expression-guided filtering to prioritize actively transcribed genes (Ward et al., 2012). This strategy kept the workflow computationally tractable while producing a draft transcriptome with more than 90% complete BUSCOs and recovering substantial expression diversity across tissues (Table 2; Figs. 3, 4). Although broader replication within each tissue could improve estimates of expression variability, our results show that sampling biologically distinct tissues and life stages can substantially expand the functional breadth represented in a probe set. Additional sampling across localities, ages, seasons, environmental conditions, and experimental treatments would likely recover further conditionally expressed genes (Rellstab et al., 2015; Isabel et al., 2020). Our approach offers a generalizable and affordable strategy for non-model species with large, complex genomes. We therefore recommend multi-tissue, expression-guided probe design as a flexible and accessible strategy for developing biologically relevant genomic tools for genomically underserved forest trees.

## Author contributions

D.S.M., E.F.G., and H.E.M. conceived and designed the study. D.S.M. conducted field and lab work and analyses. E.F.G. and J.A.K. assembled the reference transcriptome and related analyses. D.S.M. and E.F.G. wrote the manuscript. All authors contributed to the final draft. H.E.M. and W.T.P. supervised the project and acquired funding.

## Acknowledgments

The authors thank Mariel Campbell, Sandra Cervantes, Harpo Faust, Benjamin Gerstner, David Lightfoot, Mayra Macias, Michael Mann, Jessie Marlene, Melissa Sanchez, Sanna Sevanto, Lee Taylor, Dragana Vidakovic, and Christopher Witt for their support. This work was conducted as part of the University of New Mexico Women in STEM Faculty Development Fund. DSM was partially supported by NSF DBI-2305968. Computational analyses were conducted at the University of New Mexico Center for Advanced Research Computing (UNM CARC).

## Data Availability

Raw RNA-sequencing reads are available in the NCBI Sequence Read Archive under BioProject accession PRJNA1493297 (<u>link</u>). Scripts used for analyses will be made available on GitHub upon publication.

## APPENDIX S1

**Herbarium voucher, genomic resource, and sequence accession information for *Pinus edulis* RNA-seq samples.** RNA-seq reads were deposited in the NCBI Sequence Read Archive under BioProject PRJNA1493297. Tissues collected from the same adult tree share one University of New Mexico Herbarium (UNM) voucher. Residual material from all samples except the field-collected seedling was deposited in the Museum of Southwestern Biology Division of Genomic Resources (DGR); DGR catalog numbers were pending at submission. Greenhouse-grown samples were derived from collected seeds and were not herbarium-vouchered.

| **Taxon** | **Sample ID** | **Tissue type** | **UNM herbarium**<br>**voucher catalog**<br>**no.** | **DGR**<br>**catalog no.** | **SRA accession** |
| --- | --- | --- | --- | --- | --- |
| *P. edulis* | MG2-RT1-B1 | Vegetative<br>bud | UNM0155145 | Pending | SRR39573547 |
| *P. edulis* | MG2-RT1-N1 | Needle | UNM0155145 | Pending | SRR39573557 |
| *P. edulis* | MG2-RT1-P2 | Phloem | UNM0155145 | Pending | SRR39573544 |
| *P. edulis* | MG2-RT1-R2 | Root | UNM0155145 | Pending | SRR39573550 |
| *P. edulis* | MG2-RS1-N1 | Seedling<br>needle | — | — | SRR39573542 |
| *P. edulis* | MG3-RT1-B2 | Vegetative<br>bud | UNM0155144 | Pending | SRR39573546 |
| *P. edulis* | MG3-RT1-N2 | Needle | UNM0155144 | Pending | SRR39573549 |
| *P. edulis* | MG3-RT1-P2 | Phloem | UNM0155144 | Pending | SRR39573543 |
| *P. edulis* | MG3-RT1-R1 | Root | UNM0155144 | Pending | SRR39573551 |
| *P. edulis* | PO2-RT3-B1 | Vegetative<br>bud | UNM0155149 | Pending | SRR39573548 |
| *P. edulis* | PO2-RT3-N1 | Needle | UNM0155149 | Pending | SRR39573558 |
| *P. edulis* | PO2-RT3-P1 | Phloem | UNM0155149 | Pending | SRR39573545 |
| *P. edulis* | PO2-RT3-R2 | Root | UNM0155149 | Pending | SRR39573552 |
| *P. edulis* | SM-GH-S10 | Greenhouse-grown<br>seedling root | — | Pending | SRR39573556 |
| *P. edulis* | SM-GH-S16 | Greenhouse-grown<br>seedling root | — | Pending | SRR39573555 |
| *P. edulis* | SM-GH-M9 | Megagameto<br>phyte | — | Pending | SRR39573554 |
| *P. edulis* | SM-GH-M10 | Megagameto<br>phyte | — | Pending | SRR39573553 |

